# Myosin-driven fragmentation of actin filaments triggers contraction of a disordered actin network

**DOI:** 10.1101/332684

**Authors:** Kyohei Matsuda, Takuya Kobayashi, Mitsuhiro Sugawa, Yurika Koiso, Yoko Y. Toyoshima, Junichiro Yajima

## Abstract

The dynamic cytoskeletal network is responsible for cell shape changes and cell division. The actin-based motor protein myosin II drives the remodeling of a highly disordered actin-based network and enables the network to perform mechanical work such as contraction, migration and adhesion. Myosin II forms bipolar filaments that self-associate via their tail domains. Such myosin minifilaments generate both extensile and compressive forces that pull and push actin filaments, depending on the relative position of myosin and actin filaments in the network. However, it remains unclear how the mechanical properties of myosin II that rely on the energy of ATP hydrolysis spontaneously contract the disordered actin network. Here, we used a minimal *in vitro* reconstituted experimental system consisting of actin, myosin, and a cross-linking protein, to gain insights into the molecular mechanism by which myosin minifilaments organize disordered actin networks into contractile states. We found that contracted cluster size and time required for the onset of network contraction decreased as ATP concentration decreased. Contraction velocity was negatively correlated with ATP concentrations. Reduction of ATP concentration caused fragmentation of actin filaments by myosin minifilament. We also found that gelsolin, a Ca^2+^-regulated actin filament-severing protein, induced contraction of a mechanically stable network, implying that fragmentations of actin filaments in the network weaken the intra-network connectivity and trigger contraction. Our findings reveal that the disordered actin network contraction can be controlled by fragmentation of actin filaments, highlighting the molecular mechanism underlying the myosin motor-severing activities, other than the sliding tensile and compressive stress in the disordered actin network.

## Introduction

In a non-muscle cell, the actin-based cytoskeleton network, composed of apolar, disordered actin filaments associated with myosin motor proteins and actin cross-linking proteins, is responsible for a wide range of vital processes, such as cell migration and cell division. To drive these processes, the actin-based cytoskeleton network has to be reorganized, which is accompanied by actomyosin contractility (1)(2)(3). Many studies have demonstrated that actomyosin contractility mainly depends on the proper function of a motor protein, non-muscle myosin II (4)(5), which moves toward the barbed ends of actin filaments and away from the pointed ends (6). Recently, single-molecule experiments have clearly demonstrated that single molecules of non-muscle myosin II are non-processive or weakly mechanically processive (7)(8). An efficient contraction of actin-based cytoskeleton networks requires that myosin II motors, with a low duty ratio, work in assemblies of antiparallel bipolar filaments with catalytic core regions on both filament ends (hereafter referred to as myosin minifilament) (9)(10).

To date, to investigate the myosin minifilaments-driven self-organized contraction of actin-based disordered network and to develop theoretical models of the underlying molecular principles, much effort has been devoted to developing an *in vitro* minimal system consisting of purified actin, muscle myosin II, and actin cross-linking proteins (11)(12)(13). In reconstituted systems, actin filaments are bundled via cross-linking proteins. Within the bundles, myosin minifilaments and actin filaments can slide past each other. Based on the relative position of the myosin minifilament and the cross-linking protein with respect to the plus (barbed) end of an actin filament, the myosin minifilament can exert internal extensile or compressive forces, leading to extension or compression, respectively. Using such a system, several groups have demonstrated that a reconstituted actin network exhibits myosin-driven network contraction to form multiple small clusters (termed local contraction) or a single large cluster (termed global contraction), depending on the concentration of cross-linking proteins (12)(13)(14). An excess of cross-linking proteins that bundles actin filaments impedes myosin-driven actin sliding by mechanical constraints. The resultant networks are mechanically stable, whereas myosin-driven contractile stresses are maintained across it (15). Contraction of the stable network by remodeling an existing actin network requires a trigger that can remove the impediment caused by a cross-linker. Recent computational studies have suggested that buckling-induced fragmentation of actin filaments modulates network contraction (14)(16). Actin filament shortening was also demonstrated in the case of membrane-bound actin filaments (17)(18) and under extremely high concentrations of myosin (19). However, *in vitro* experiments have not shown clearly how myosin minifilament-driven fragmentation of actin filaments regulates contraction of the three-dimensional (3D) actomyosin disordered networks.

## Materials and Methods Proteins

Actin and myosin were purified from the rabbit skeletal muscle (20). Actin was stored in a G buffer [2 mM Tris-HCl (pH 8.0), 0.2 mM ATP, 0.2 mM CaCl_2_, and 0.2 mM DTT] in liquid nitrogen. Myosin was stored in 600 mM KCl and 1 mM DTT in liquid nitrogen. Recombinant human anillin was prepared as follows. First, cording sequence of the multi-functional GFP (21) was attached to the 3’ end of the human anillin cDNA using F-primer (5’-GGG AAG CTT GCC ACC ATG GAT CCG TTT ACG GAG AAA C -3’) and R-primer (5’-GGG ACG CGT AGG CTT TCC AAT TGG TTT GTA G - 3’). The construct encoding anillin-mfGFP was cloned into pcDNA5/FRT/TO expression vector (Invitrogen) for HEK293F cell expression. The transfection and purification methods were as described elswhere (21)(22). Anillin-mfGFP was stored in an anillin buffer [10 mM PIPES-Na (pH 7.0), 150 mM NaCl, 1 mM DTT, and 10% sucrose (w/v)] in liquid nitrogen. Recombinant human gelsolin was prepared as follows. First, a hexa histidine-tag was attached to the 3’ end of human gelsolin DNA sequence (23). The gelsolin-His construct was cloned into pCold III vector for expression in *Escherichia coli* BL21 Star (DE3) cells. The pCold-gelsolin-His were next transformed into the cell. Gelsolin-His was expressed for 24 h at 15°C; the cells were collected by centrifugation and resuspended in a lysate buffer (20 mM imidazole-HCl, 300 mM KCl, 1 mM DTT, and 10 mM CaCl_2_) containing the protease inhibitor cocktail (Roche), and sonicated for 20 min on ice. Next, the lysate was centrifuged for 20 min at 305,000 × g and at 4°C. Ni-NTA resin (Bio-Rad) was added and the mixture gently stirred for 1 h at 4°C. The Ni-NTA beads were washed with a buffer (20 mM imidazole-HCl, 300 mM KCl, 1 mM DTT and 10 mM CaCl_2_) by centrifuging several times. Gelsolin-His was then eluted with another buffer (200 mM imidazole-HCl, 300 mM KCl, 1 mM DTT and 10 mM CaCl_2_), desalted on a NAP5-culumn (GE), and stored in a gelsolin buffer [10 mM Tris-HCl (pH 7.4), 10 mM CaCl_2_, 150 mM NaCl, 1mM DTT and 50 μM ATP] at liquid nitrogen.

The concentrations of actin, meromyosin, and myosin were determined based on ultraviolet absorbance of their solution, using the absorption coefficients 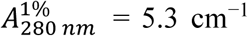 for full-length myosin II, 6.0 cm^-1^ or HMM and 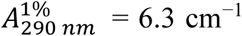 for G-actin, and using molecular masses of 470 kDa for full-length myosin II, 350 kDa for heavy-meromyosin (HMM) and 42 kDa for actin (24). The concentrations of anillin and gelsolin were estimated by SDS-PAGE on 10% acrylamide gels using BSA standards (Thermo Scientific) loaded on the same gel. Gels were stained with Quick-CBB PLUS (Wako) and imaged using a CCD camera (CSFX36BC3, Toshiba). The bands containing anillin, gelsolin, and BSA standards were quantified using ImageJ (NIH).

## Microscopy

Actin filament was observed using a fluorescence microscope (IX70, Olympus) with a stable stage (KS-N, ChuukoshaSeisakujo) and a stage controller (QT-CM2-35, Chuo Precision Industrial); with illumination from a mercury lump (U-HGLGPS, Olympus); 20 ×/ NA, UPlanFl objective lens (Olympus); and U-MWIG filter set (Olympus). Images were recorded by sCMOS camera (Zyla 4.2 PLUS USB3.0, Andor). Shutter was controlled using SSH-C (Sigma-koki). In the actin fragmentation assay, 100×/ NA 1.40 oil immersion UPlanApo objective lens (Olympus) was used.

## Polymerization of myosin minifilament

Myosin stored in the stock buffer containing 600 mM KCl and 1 mM DTT was diluted in the assay buffer [50 mM PIPES-Na (pH 7.4), 50 mM KCl, 1 mM CaCl_2_, 1 mM MgCl_2_, and 1 mM EGTA] to form myosin minifilament at room temperature (24 ± 1 °C). The final buffer contained 33 mM PIPES-Na, 233 mM KCl, 0.66 mM CaCl_2_, 0.66 mM MgCl_2_ and 0.66 mM EGTA.

## Myosin minifilaments-driven network contraction assay

Actomyosin network formation was initiated by mixing myosin minifilaments, anillin and G-actin in an assay buffer [50 mM PIPES-Na (pH 7.4), 50 mM KCl, 1 mM CaCl_2_, 1 mM MgCl_2_, and 1 mM EGTA] containing 2 mg/ml BSA, 1.0 μM rhodamine phalloidin, 0.5 mM DTT, and varying concentrations of ATP. To prevent photobleaching, 0.2% catalase, 1.5 mg/ml glucose, and 50 U/ml glucose oxidase as the oxygen scavenger were added (23). For a constant ATP concentration in an assay, 20 mM creatine phosphate and 0.3 mg/ml creatine kinase were added to the solution as the ATP-regenerating system (23). The assay was performed in flow chambers assembled from two coverslips, 18 × 18 mm and 24 × 36 mm (Matsunami), attached with a double-sided tape (NW-25, Nichiban). The glass surface was blocked with 2 mg/ml BSA to prevent glass-protein interactions. The assay mixture was placed in the chamber, and the observation commenced 30 s after mixing. For the assay, both ends of the chamber were sealed with nail polish. Images were recorded by sCMOS camera every 10 s, with 0.5-s exposure time. Once the image acquisition was complete, the entire chamber area was inspected to identify the largest cluster.

Network contraction velocity (CV) and contraction initiation time (CIT) were calculated as follows. First, background intensity was subtracted using a built-in “subtract background” plugin in ImageJ; all time-lapse images were transformed using the same low limit threshold. Next, the network area in each frame was determined. The CV was calculated as the slope of the decreasing network area as a function of time (in s). The CIT was the time when the measured area began to decrease. The maximum cluster size was calculated as follows. First, background intensity was subtracted using a built-in “subtract background” plugin in ImageJ. To determine the cluster size, a built-in “analyze particle” plugin was used. Maximum cluster size was the size of the largest cluster size in the final image, in the entire glass chamber area.

## Direct observation of severed F-actin in a three-dimensional (3D) disordered network

F-actin was prepared as follows. First, G-actin was polymerized in an F buffer [5 mM Tris-HCl (pH8.0), 1 mM MgCl_2_, 100 mM KCl, 0.1 mM CaCl_2_, and 0.5 mM DTT] containing 10 μM rhodamine phalloidin, and 100 μM ATP, for 1 h at room temperature. The mixture was then centrifuged for 1 h at 355,000 × g and 4 °C (Optima MAX, Beckman), and the pellet was suspended in the F buffer containing 1 μM ATP. To observe F-actin in a 3D disordered network, partly labeled actin network was prepared, containing 3 μM G-actin, 1 μM dark phalloidin, 0.1% rhodamine labeled F-actin, 0.36 μM anillin-GFP, 1 μM myosin minifilaments, the oxygen scavenger, 2 mg/ml BSA, and 1,000 μM ATP in the assay buffer. The images were recorded by sCMOS camera every 2 s, with 0.5-s exposure time. To calculate the velocity of the F-actin fragment, the fragment’s movement was quantified using an automated tracking program Mark2 (Ken’ya Furuta, NICT). Fluorescence images of the fragment were fitted using a 2D Gaussian function to determine the position of the fluorescence intensity peak, corresponding to the center of the fragment (25). Moving distance of the actin fragment was calculated using the (x, y) coordinate of the fragment in each frame. Fragment velocity was calculated based on the moving distance during a 10-s time interval (five frames).

## Fragmentation of F-actin bound to glass surface coated with a non-ionic surfactant

The glass chamber was coated with the non-ionic surfactant F127 as follows. The cover glass was first washed in 5 M KOH for 12 h, and thoroughly washed in MilliQ water. The KOH-cleaned glass was next silanized using silanization solution I (SIGMA-ALDRICH), and washed with acetone (twice) and MilliQ water (twice). The chamber was assembled from two coverslips, 18 × 18 mm and 24 × 36 mm, of silanized glass, attached with a double-sided tape (NW-25, Nichiban); 25 μg/ml of anti-GFP antibody (Takara) was placed in the chamber. After 3 min, the chamber was washed three times with the assay buffer, and F127 was added. After 3 min, unbound F127 was washed away with the assay buffer, three times. Then, 0.1 μM anillin-GFP was loaded into the glass chamber coated with the non-ionic surfactant. After 3 min, chamber was washed twice with the assay buffer. Next, 0.1 μM solution of rhodamine phalloidin-labeled actin filaments was added and immobilized on the glass surface via anillin-GFP. Finally, 1 μM myosin minifilament solution with different ATP concentrations was added. The solution contained 0.3% methylcellulose to localize F-actin to the glass surface, ATP-regenerating system, and the oxygen scavenger in the assay buffer. Images were recorded by sCMOS camera every 1 min, with 0.5-s exposure time. The length of actin filaments was calculated by using the built-in “segmented-line tool” in ImageJ. The lower measurement limit of the actin filament length was considered to be ∼0.4 μm.

## Gelsolin-induced network contraction assay

To sever actin filaments in a prestressed network, 1 μl of 0.02-0.20 μM gelsolin diluted in the gelsolin buffer [10 mM Tris-HCl (pH 7.4), 10 mM CaCl_2_, 150 mM NaCl, 1mM DTT and 50 μM ATP] was loaded into the glass chamber. The final concentration of gelsolin was 0.002-0.020 μM. Images were recorded by sCMOS camera every 10 s, with 0.5-s exposure time.

## Measurement of myosin minifilament length using electron microscopy

A 10-nM minifilament solution was applied onto carbon-coated copper grids and negatively stained with 1.0% (w/v) uranyl acetate (26). The specimens were examined using an electron microscope (Hitachi, H-7500) at magnification of 12,000 ×, operated at 80 kV. The length of myosin minifilaments was evaluated visually by using the built-in “segmented-line” tool in ImageJ.

## Results

### Various contraction of a 3D disordered network of actomyosin and anillin

To explore the mechanism by which a disordered actomyosin network contracts, we performed an *in vitro* reconstituted myosin minifilament-driven contractility assay (**Fig. 1 *A***). In the assay, a fixed concentration of monomeric actin (G-actin, 3 μM), and varying concentrations of myosin II and actin cross-linking protein (ACP) were mixed; constructions of the 3D disordered actin networks were then observed using fluorescence microscopy. During network construction, actin monomers assembled spontaneously into actin filaments (F-actin), which was stabilized by rhodamine phalloidin. To cross-link the resulting actin filaments, we used anillin, an ACP that is localized at the contractile ring during cytokinesis (27). In the presence of 0.36 or 0.72 μM anillin, networks were a homogenous mesh of actin filaments; at 0.90 μM anillin, bundled, mesh-like actin networks cross-linked by anillin were formed (see **Fig. S1** and **Movie 1 in Supporting Material**). To create internal tensile and compressive stresses within the actin networks cross-linked by passive ACP, the motor activity of myosin II was used (28)(**Fig. 1 *A***). Dimeric myosin II purified from the rabbit skeletal muscle self-assemble through an interaction of their C-terminal antiparallel coiled-coil domains into bipolar myosin minifilaments with motor domains at each end (29). Bipolarity of the myosin minifilament allows actin filaments of opposite orientations to slide toward each other (30)(31). In the assay, myosin minifilaments were ∼0.65 μm long, based on electron microscopy analysis (see **Fig. S2**), i.e. approximately 100 myosin molecules per filament (9). When a fixed overall myosin concentration (1 μM) was used in a series of actin networks reconstituted with varying concentrations of anillin in the absence of an ATP-regenerating system, distinct self-organizing structures of different lengths were reproducibly observed (**Fig. 1 *B***), which was consistent with previous experimental (13) and theoretical (14)(15) studies. In the absence of anillin, the myosin minifilaments were still able to contract the actin network and form innumerable small clusters with the mean maximum cluster size below 100 μm^2^ (hereafter referred to as small clusters; **Fig. 1 *B***, left-most column, **Fig. 1 *C***, open circle and see **Movie 2 in Supporting Material**). These clusters often combined, slightly increasing the overall cluster size (32). This indicated that myosin minifilaments not only generate contractile force but also cross-link actin filaments, as predicted by theoretical studies (13)(15). With increasing anillin concentration (0.36 μM), myosin activity broke the passive cross-linked actin network into multiple disjointed medium-size clusters with a mean maximum cluster size of 100 to 1,000 μm^2^ (hereafter referred to as medium clusters; **Fig. 1 *B***, second column from the left, **Fig. 1 *C***, open triangle and see **Movie 3 in Supporting Material**). When a larger amount of anillin (0.72 μM) was added, a highly cross-linked actin network was contracted with a single dense large cluster with a mean maximum cluster size of over 1,000 μm^2^ (hereafter referred to as the large cluster; **Fig. 1 *B***, third column from the left, **Fig. 1 *C***, open diamond and see **Movie 4 in Supporting Material**). Typically, self-organized structures reached a steady state within 1 h of mixing, depending on the cluster size (hereafter referred to as CS). Thus, myosin activities led to spontaneous formation of various contracted structures, which was not observed in the corresponding passive cross-linked actin networks in the absence of myosin. However, at a still higher concentration of anillin (0.90 μM), excessively bundled and cross-linked actin networks were formed, but no contraction was observed (hereafter referred to as the prestressed network; the-right most column in **Fig. 1 *B*** and **Fig. 1 *C***, open square, and see **Movie 5 in Supporting Material**) (15). Further, we controlled the internal stresses by varying the amount of myosin II in the presence of fixed concentrations of G-actin and anillin (3 μM and 0.36 μM, respectively), and observed the self-organizing process of the disordered actin network. With gradually increasing myosin concentrations, the networks contracted on a larger CS (**see Fig. S3**). Our results, with anillin as the cross-linker, confirmed previous findings and theoretical predictions that network connectivity depends on the amount of both the cross-linking proteins and motor proteins, and regulates the CS (13)(14)(15).

**Figure 1:**
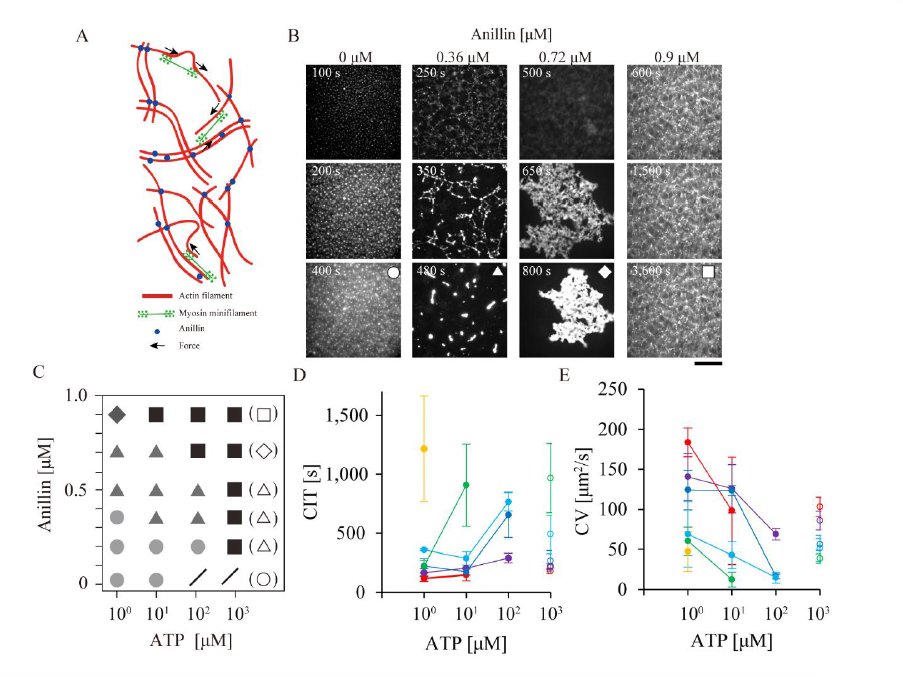
Contraction of a 3D disordered network of actomyosin and anillin. (*A*) Diagram of an active actomyosin network with the cross-linker, anillin. Actin filaments (red lines) are cross-linked by anillin (blue dots). Myosin minifilaments (green) drive actin compaction (black arrows) in the 3D disordered network. (*B*) Cluster size (CS) depends on anillin concentration. Time-lapse images of network contraction in the presence of various anillin concentrations are shown. Concentrations of G-actin (3 μM) and myosin (1 μM) were constant. The initial ATP concentration was 1000 μM, and it decreased with time. Scale bar, 100 μm. Based on CS, clusters are categorized as small (circle), medium (triangle), or large (diamond). (*C*) Phase diagram of CS depends on the concentrations of both ATP and anillin. Open and filled symbols indicate the CS in the absence and presence of the ATP-regenerating system, respectively. The symbols are the same as (*B*). When the network does not break up and, therefore, does not form clusters, the state is defined as prestressed network (square). Slash sign indicates no specific actin-based structures. (*D*) Contraction initiation time (CIT) at different ATP concentrations. All networks contain various concentrations of ATP with the ATP-regenerating system. CIT is defined as the time required for the onset of network contraction. With decreasing ATP concentration, CIT become shorter. With increasing anillin concentration, CIT become longer. Filled and open dots indicate CS in the presence and absence of the ATP-regenerating system, respectively. Anillin was added at 0 μ M (red), 0.2 μ M (purple), 0.36 μ M (blue), 0.5 μ M (sky blue), 0.72 μ M (green) or 0.9 μ M (yellow). (*E*) Contraction velocity (CV) at different ATP concentrations. With decreasing ATP concentration, CV increased. This is inconsistent with the myosin-driven actin filament sliding velocity at low ATP concentrations in a classical gliding assay (33). The colors are the same as (*D*).

## ATP-dependent contractility in a 3D disordered actomyosin network

To probe the roles of the mechano-chemical activity of myosin-minifilament in the contraction mechanism, we quantified the ATP-dependent CS (**Fig. 1 *C***). When a fixed myosin minifilament concentration (1 μM) was used in a series of networks with varying concentrations of anillin and ATP in the presence of an ATP-regenerating system, contractions only occurred at sufficiently low ATP concentrations (**Fig. 1 *C***) (32)(34), indicating that ATP concentration also affected CS. At high ATP concentrations, the contractions were incomplete, and the actin network was similar to the network formed by only actin and passive cross-linker anillin (see **Fig. S1**). We also found that the time required for the onset of a major structural change in the network, i.e., the contraction initiation time (hereafter referred to as CIT), changed in both ATP and anillin concentration-dependent manners (**Fig. 1 *D***). The myosin minifilament-driven contraction velocity (hereafter referred to as CV) also depended on both the ATP and anillin concentrations (**Fig. 1 *E***). Intriguingly, as ATP concentration decreased, CIT shortened and CV increased. Since myosin-driven actin filament sliding in general slows down as ATP concentration decreases (33), myosin minifilament-driven contraction should be regulated by an ATP-dependent activity of the myosin minifilament other than motility.

## Myosin minifilament-induced fragmentation of actin filaments in a 3D disordered network

To understand what determines the CIT and CV parameters, we first analyzed individual actin filaments during contraction by introducing a small amount of rhodamine phalloidin-labeled actin filament (0.1%) into a 3D disordered network of unlabeled 3 μM G-actin, 1 μM myosin, 0.5 μM anillin, and 1 mM ATP in the absence of ATP-regenerating system (**Fig. 2 *A*** and ***B*** inset and see also **Movie 6 in Supporting Material**). We found that the labeled actin filaments slid, and the filaments were frequently fragmented in the network. The most actin filament segments moved to assemble in small clusters. We also tracked the myosin minifilament-driven sliding of actin filament segments to investigate the relationship between the moving velocity and actin filament fragmentation during network contraction (**Fig. 2 *B***). The moving velocity fluctuated, whereby a velocity increase was often followed by actin filament fragmentation. Approximately 70% of fragmented actin filaments immediately after filament severance moved twice as fast as actin filaments before severance. These results implied that actin filament breakage resulted in a release of drag forces generated by anillin cross-linked to other non-fluorescent (i.e., experimentally invisible) actin filaments.

**Figure 2:**
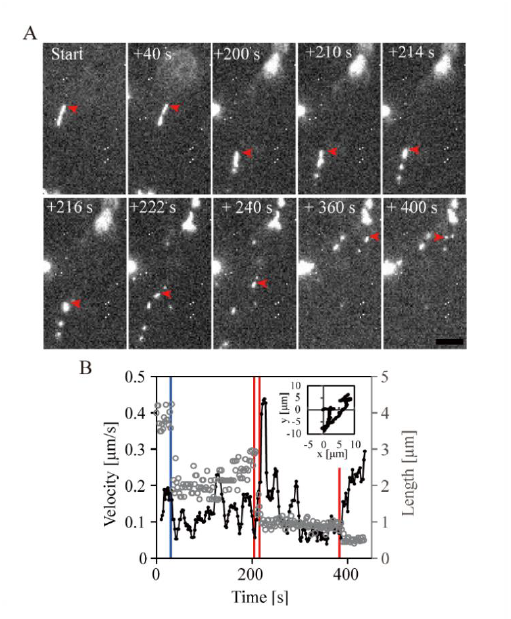
Myosin minifilament-induced fragmentation of actin filaments occurs in a 3D disordered network. (*A*) Time-lapse images of an actin filament fragmentation in the 3D disordered network formed from 3 μM G-actin, 1 μM phalloidin, 1 μM myosin, 0.5 μM anillin, 1000 μM ATP, and 0.1% rhodamine phalloidin-labeled actin filaments. Time (in s) after the initiation of observation is shown. The observation commenced 6 min after mixing. Red arrow-heads point to the measured fragment. Scale bar 5 μm. (*B*) The moving velocity of a fragmented actin filament (black dots and line) pointed out by red arrow heads in **Fig. 2 *A***, and the length of fragmented actin filament (gray circles) as a function of time. Red lines denote the fragmentation of actin filament concomitant with an acceleration event of actin filament movement (212 s, 218 s, and 390 s). Blue line denotes fragmentation of actin filament without an acceleration event (40 s). Acceleration events with fragmentation occurred at a rate of 70% (N = 10 experiments). Inset, trajectory of severed actin filament.

## Fragmentation of actin filaments driven by myosin-minifilaments at lower ATP concentrations

To resolve the nature of the interplay between fragmentation of actin filament and ATP-dependent CIT and CV, we quantified the length of actin filament segments with varying ATP concentrations (**Fig. 3**). Fragmentation of actin filaments by myosin minifilaments from the skeletal muscle (the same as ones used in the current study) was previously demonstrated in a 2D-quasi actin network consisting of actin filaments tethered to supported lipid bilayers; the fragmentation occurred only at ATP concentrations of 0.3 and 1.0 μM (17). In the current study, rhodamine phalloidin-labeled actin filaments were immobilized on an anillin-coated glass surface; then, myosin minifilaments (fixed concentration) and ATP (varying concentrations) were added (**Fig. 3 *A***). In the presence of 1 mM ATP, the length of actin filaments did not change substantially (**Fig. 3 *B*** and ***D***). However, with reduced ATP concentrations, the average actin filament length was reduced, indicating a negative correlation between actin filament severing activity and ATP concentration (**Fig. 3 *C*** and ***D***). A control experiment demonstrated that the addition of heavy meromyosin (HMM) did not show fragmentation of actin filaments, and that the observed fragmentation required the activity of myosin minifilaments (see **Fig. S5**). This finding, together with the positive correlation between ATP concentration and CIT (**Fig. 1 *D***), and the negative correlation between ATP concentration and CV (**Fig. 1 *E***), led to the idea that fragmentations of actin filaments in the actin network promote the network contraction. Although our results clearly showed that myosin minifilaments alone were able to sever actin filaments and that low ATP concentrations promoted the myosin minifilament-driven severing activity, there remains the possibility that other properties of the myosin minifilament, such as high-processivity or generation of a strong sliding force (17)(35), associated with low ATP concentrations are essential for the CIT and CV.

## Network contraction induced by actin-severing protein

We next asked if fragmentation of actin filaments could constitute one of the origins of contraction, breaking the mechanical stability of the networks. To this end, we let the prestressed network to form using 3 μM G-actin, 1 μM myosin, 0.72 μM anillin, and 50 μM ATP in the presence of the ATP-regenerating system. Then, we added gelsolin, an actin filament-severing protein, and examined the occurrence of contraction. Following the addition of gelsolin, the prestressed network began to contract locally, and the contraction propagated to the whole network, resulting in multiple medium-size clusters (**Fig. 4 *A*** and see also **Movie 7, left panel**). A control experiment demonstrated that the addition of a buffer solution without gelsolin did not disturb the prestressed network, and that the observed contraction was not an artifact of mechanical perturbation (**Fig. 4 *B*** and see also **Movie 7, right panel, in Supporting Material**). We also found that the CV relied on gelsolin concentration (**Fig. 4 *C***). These results indicated that fragmentation of actin filaments by gelsolin perturbs the force balance between the drag force generated by anillin cross-linking and the compressive force generated by myosin power-stroke bound to actin filament, resulting in contraction of the prestressed network.

**Figure 3:**
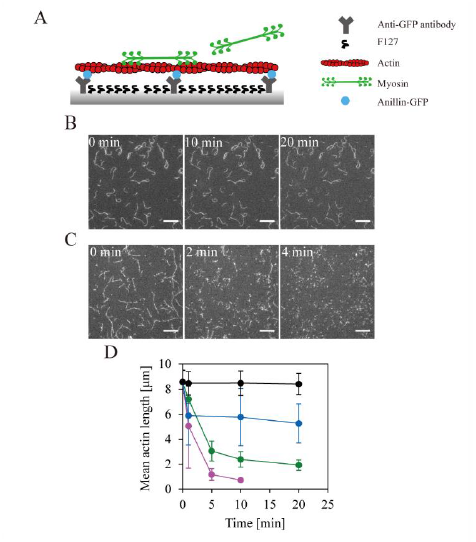
Fragmentation of actin immobilized on a glass surface coated with a non-ionic surfactant. (*A*) Schematic illustration of the experiment. (*B*) and (*C*): Fluorescence images of rhodamine phalloidin-labeled actin filament fragmentation in the presence of 1000 μM ATP (*B*) or 1 μM ATP (*C*), and an ATP-regenerating system. Scale bar, 10 μm. (*D*) ATP concentration dependency of actin fragmentation. Mean actin filament lengths in the presence of 1–1000 μM ATP as a function of time. Actin filaments were immobilized on the glass surface via an anillin-GFP construct and anti-GFP antibody. Myosin minifilaments severed and shortened actin filaments over time. The final mean lengths (μm) of actin fragments were 0.7 ± 0.2 (mean ± S.D), in the presence of 1 μM ATP (magenta; N_exp_ = 3; the total number of actin filaments analyzed during N_exp_ independent experiments: N_fil_ = 236); 1.9 ± 0.4, in the presence of 10 μM ATP (green; N_exp_ = 3, N_fil_ = 471); 5.3 ± 1.6 in the presence of 100 μM ATP (blue; N_exp_ = 3, N_fil_ = 229); and 8.4 ± 0.9, in the presence of 1000 μM ATP (black, N_exp_ = 3, N_fil_ = 218).

**Figure 4:**
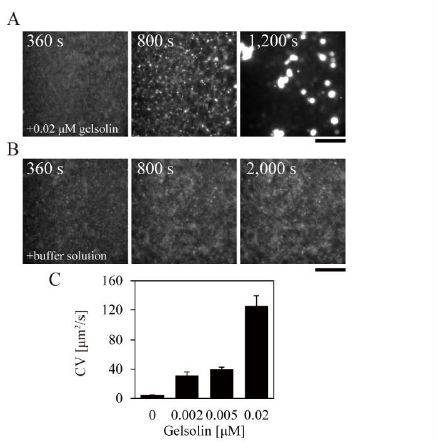
The actin-severing protein, gelsolin, induces network contraction. Sequential images of a disordered network are shown. The network comprised 3 μM G-actin, 1 μM rhodamine phalloidin, 1 μM myosin, 0.72 μM anillin, 50 μM ATP, and the ATP-regenerating system. Gelsolin (*A*) or buffer containing 50 μM ATP (*B*) was additionally injected. Scale bar, 100 μm. (*A*) When a small amount of gelsolin is added (360 s) to the stable 3D disordered network, the network contracts to form medium clusters (1200 s). (*B*) When a buffer containing 50 μM ATP is added (360 s) to the stable 3D disordered network, contraction does not occur (2000 s). (*C*) Changes in the CV with varying gelsolin concentrations: 0 μM (N_exp_ = 3), 0.002 μM (N_exp_ = 3), 0.005 μM (N_exp_ = 4), and 0.02 μM (N_exp_ = 4). The data are presented as the mean ± S.E.; N_exp_, number of independent experiments.

## Discussion

Actin-based cytoskeletal network contraction requires myosin activity driven by ATP hydrolysis (36). The contraction did not occur in the presence of a nonhydrolyzable ATP analog 5’-adenylyl-imidodiphosphate (AMPPNP) (**Fig. S4, upper panel**) or the myosin II ATPase inhibitor blebbistatin (**Fig. S4, middle panel**) (5). Myosin II is a non-processive motor with a low duty ratio, implying that these motor molecules need to function together, in an ensemble, to produce a substantial motion and force. Non-processive HMM that does not form minifilaments does not contract the network within a range of 1–1,000 μM ATP (**Fig. S4, lower panel**), while myosin minifilaments drive contractile forces in actin networks (**Fig. 1 *C***). At ATP concentrations below an apparent *K*_*m*_ (∼0.2 mM) for myosin II-driven actin sliding (33), where each myosin motor consisting of the minifilament was able to spend a longer period attached to actin filament during each ATP hydrolysis cycle, network contraction was observed. In contrast, at ATP concentrations exceeding the apparent *K*_*m*_, network contraction on the cellular scale was not observed, probably because myosin minifilaments became weakly processive for their low duty ratio at higher ATP concentrations (17), and did not produce enough force to remodel the actin network.

Numerous studies have also shown that ACP is a major contributor to network contraction (13)(37)(38). Cross-linking of actin filaments by ACP imposes isotropic dot-like constraints on the network. When the ACP concentration is increased gradually, the cross-linked networks become stiffer (39). Our study revealed that, in proportion to network connectivity regulated by the amount of anillin, as reported for different ACPs (12)(13)(14), myosin minifilament-driven CS increased (**Fig. 1 *B*** and ***C***), CIT became longer (**Fig. 1 *D***), and CV slowed (**Fig. 1 *E***). This implied that myosin-driven contraction force was inhibited by drag force exerted by anillin bound to actin filaments. At a higher concentration of ACP, under conditions leading to a much stiffer prestressed network, stall force generated by myosin minifilaments was balanced with drag force generated by anillin cross-linking. Even such prestressed networks could be contracted at sufficiently low ATP concentrations (**Fig. 1 *D***). Interestingly, CS decreased as the ATP concentration decreased. The finding is counterintuitive; connectivity in the actomyosin network increases at low ATP concentration, when myosin minifilament becomes a high duty ratio motor.

What weakens the connectivity at low ATP concentrations? One clue was provided by the *in vitro* fragmentation assay, which revealed that at low ATP concentrations the occurrence of myosin minifilament-driven fragmentation of actin filaments bound to a glass surface was likely (**Fig. 3**). The most obvious explanation for this is that the actin filament severance occurred in the bare zone within the minifilament. Buckling-induced breaking force of actin filament in the bare zone was simulated to be ∼20 pN (17), which might be achievable by the myosin minifilament. Within the network, such severance of actin filaments induces cracks in the network (14), resulting in weak connectivity. In fact, direct observation of actin filaments in actin networks using fluorescence microscopy demonstrated that actin filaments were often fragmented, and the moving velocity of the actin filament segment immediately after filament severance often increased (**Fig. 2**). Gelsolin-driven fragmentation of actin filaments also caused prestressed networks to contract with medium clusters (**Fig. 4**). Thus, the fragmentation of actin filament functions to reduce network connectivity and triggers contraction.

How does fragmentation promote contraction? In disordered actin networks, mechanical constraints of actin filaments generated by ACP cross-linking act to impede myosin minifilament-driven sliding of actin filaments. Single myosin motor molecules generate stall forces of several pN, and myosin minifilament-driven actin filament sliding ceases with the overwhelming load imposed by ACP cross-linking (40). The cross-linked actin bundles locally experience tension, and the prestressed network is mechanically stiffened and appears homogeneous on a macroscopic scale (**Fig. 5 *i***). When myosin minifilaments sever the actin filaments, the mechanical rigidity of the actin networks decreases locally. Myosin minifilament-driven fragmentations of actin filaments occur everywhere in the highly cross-linked network, which in turn induces the intra-network connectivity to be weakened, and thence stress is redistributed (**Fig. 5 *ii***). This allows local myosin-driven sliding of actin filaments from the area where the symmetry induced by equivalent contractile force and cross-linking constraints would collapse promoting network deformation. A previous *in vitro* sliding assay using myosin minifilaments showed that the velocity pulling actin filaments toward the central, myosin head-free bare zone is about nine-times faster than that pushing actin filaments away from the bare zone (30). Superiority of the actin filament pulling velocity to the actin filament pushing velocity may cause compaction of actin filaments, resulting in contractile deformations (**Fig. 5 *iii***). Two actin filaments sliding over each other, driven by a myosin minifilament, leads to the formation of actin filament overlaps that can be bundled by anillin (38). More extensively cross-linked bundles inhibit the sliding actin filament from being extended (**Fig. 5 *iv***).

**Figure 5:**
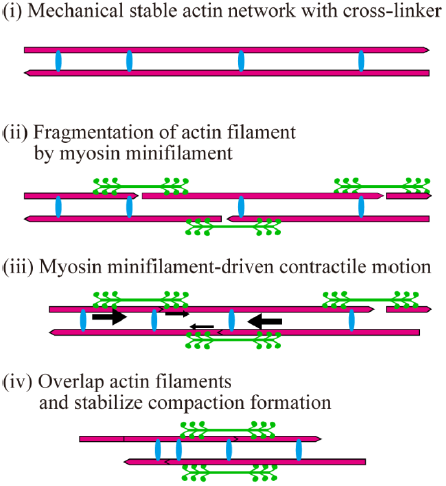
Proposed model explaining how myosin minifilament-driven actin fragmentation triggers the contraction of a mechanically stable disordered actin network. Actin filaments, magenta line; anillin, blue dots; myosin minifilaments, green shapes. The thickness of black arrows indicates the relative velocity of sliding actin fragment. Upon actin filament fragmentation, the mechanical stability network is weakened, and the network local contraction propagates the large-scale contractions (see text for details).

In conclusion, myosin minifilament-driven severance of the actin filament is a key factor in the timing of network contraction. The presented model in which actin fragmentation triggers contraction *in vitro* in a 3D disordered network may be also applied *in vivo*. In a cell, the structure of actin networks changes constantly, and this is controlled by various proteins, such as several non-muscle myosin isoforms (41)(42), passive cross-linkers (43), capping proteins (44), and severing proteins such as cofilin (45) and gelsolin(46). These constitute an indispensable regulating mechanism of such networks as the filopodium and lamellipodium (47). To our knowledge, actin fragmentation by myosin minifilament *in vivo* has not been reported. However, it is plausible because of the existence of distinct types of non-myosin II motors with different duty ratios (41). E.g., non-muscle myosin IIB involved in contractile ring contraction is a high-processive motor with a high duty ratio (7). A mechanically stable contractile ring might start to contract upon fragmentation of actin filaments by non-muscle myosin IIB. It would be interesting to evaluate whether non-muscle myosin IIB exhibits severing activity in the presence of high ATP concentrations, and observe myosin-driven fragmentation *in vivo*.

## SUPPORTING MATERIAL

Five figures, seven movies are available at http://www.biophsj.org/biophysj/supplemental/.

## Acknowledgments

We thank R. Uehara for providing human cDNA clones. This work was supported in by JSPS KAKENHI (No.15H01629, No.15K07022 No.15H02006, No.16KT0065 to J.Y.). The authors declare that they have no competing financial interests.

## Author Contributions

J.Y. designed the project. K.M. performed all experiments and analyzed all data. T.K. constructed and purified anillin. S.M. set up imaging system. T.K. and Y.K. collected EM data. Y.Y.T purified actin and myosin. All the authors wrote the manuscript.

## SUPPORTING MATERIAL

**Figure S1:**
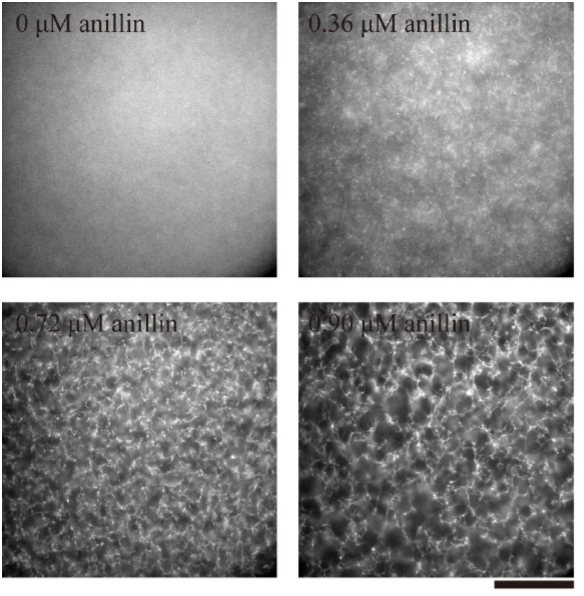
Dependence of actin bundle networks on anillin concentration in the absence of myosin. The following reagent concentrations were used in the depicted experiments: 3 μM G-actin, 1 μM rhodamine phalloidin, 1 mM ATP (without ATP regenerating system), and 0-0.9 μM anillin. Scale bar, 100 μm. In the absence of anillin, no structure of actin network was observed. In the presence of 0.36 or 0.72 μM anillin, actin networks were a homogenous mesh of actin filaments. In the presence of 0.90 μM anillin, bundled, mesh-like actin networks cross-linked by anillin were formed.

**Figure S2:**
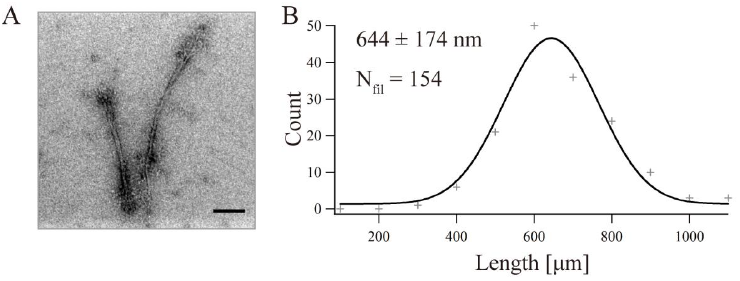
Length distribution of myosin minifilaments. (A) Electron microscopy images of negatively stained myosin minifilaments. Two myosin minifilaments are shown. Scale bar, 100 nm. (B) Length distribution of myosin minifilaments. The mean actin filament length and standard deviation were derived from a Gaussian fitting. N_fil_ is the total number of filaments.

**Figure S3:**
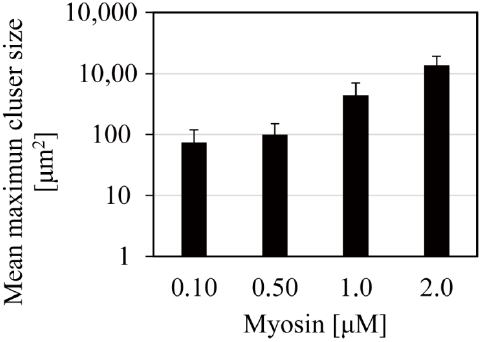
Mean maximum cluster size at different myosin concentrations. In the assay, 3 μM G-actin, 0.36 μM anillin-GFP and 0.1-2.0 μM myosin minifilament were mixed in assay buffer containing 1 μM rhodamine phalloidin, 2 mg/ml BSA and 1 mM ATP. At the myosin concentration of 0.1, 0.5, 1.0, or 2.0 μM, the mean maximum cluster size (μm^2^) was 744 ± 46 (mean ± S.E.; the number of independent experiments; N_exp_ = 4), 99 ± 51 (N_exp_ = 4), 440 ± 265 (N_exp_ = 4) and 1369 ± 558 (N_exp_ =5), respectively.

**Figure S4:**
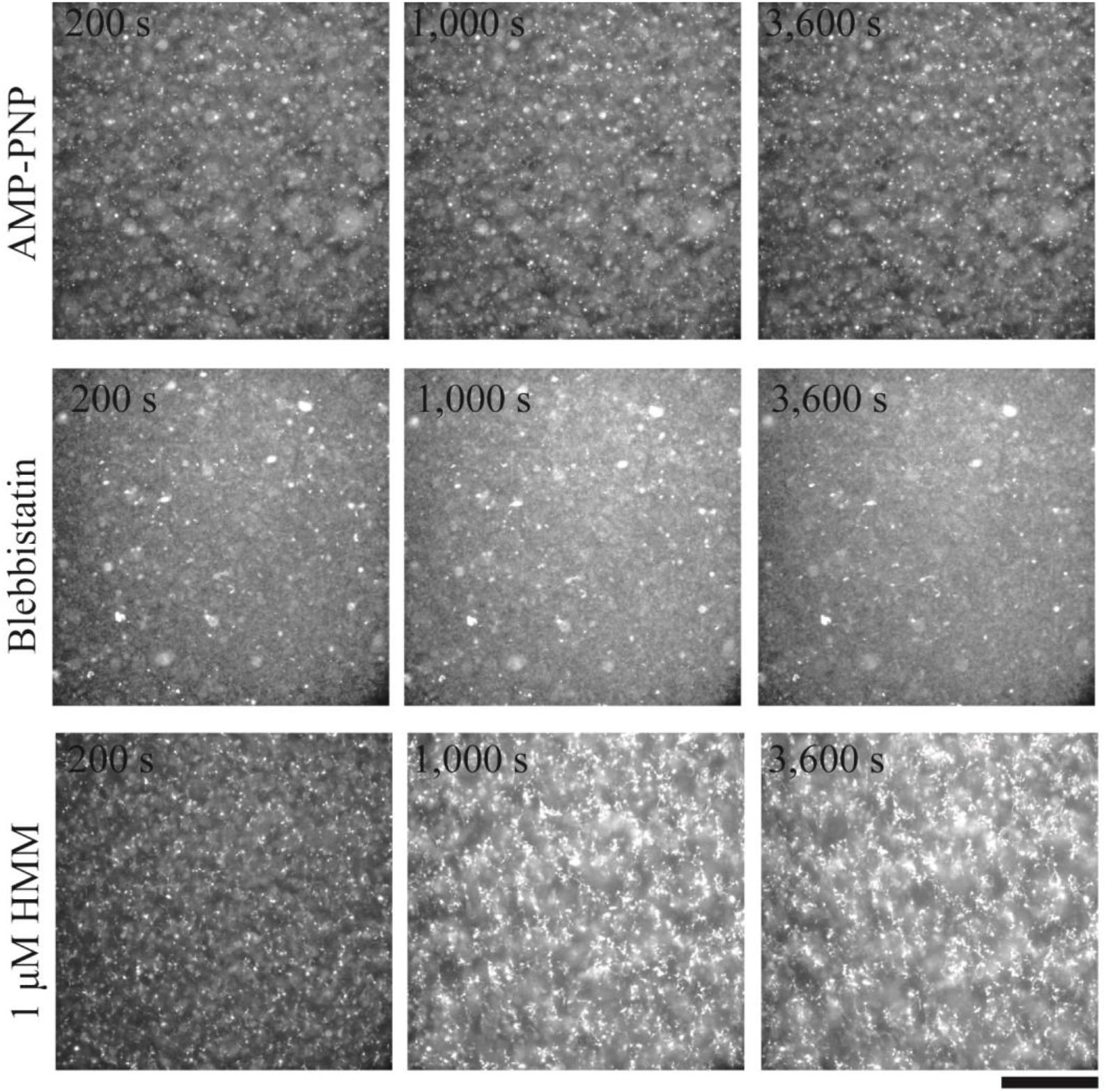
Actin-based cytoskeletal network contraction requires myosin activity driven by ATP hydrolysis. (Upper panel) The structure of a network containing 3 μM G-actin, 1 μM myosin, 0.36 μM anillin, 1 μM rhodamine phalloidin and 1 mM AMP-PNP remained unchanged. (Middle panel) The structure of a network containing 1 μM blebbistatin, 3 μM G-actin, 1 μM myosin, 0.36 μM anillin, 1 μM rhodamine phalloidin and 1 mM ATP remained unchanged. (Lower panel) A network containing 3 μM G-actin, 1 μM HMM, 0.36 μM anillin, 1 μM rhodamine phalloidin and 1 mM ATP showed a slight movement but the structure remained largely unchanged.

**Figure S5:**
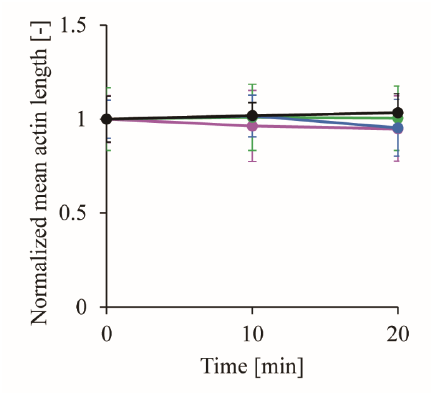
HMM did not show actin filament severing activity. Normalized mean actin lengths in the presence of 1μ M (magenta), 10 μ M (green), 100 μ M (blue) and 1000 μM (black) ATP with ATP regenerating system as a function of time. Actin filaments were immobilized on the glass surface via an anillin-GFP construct and anti-GFP antibody. 3 μ M G-actin and 1 μ M HMM were used. Concentrations of G-actin and myosin motor heads are the same as Figure 3.

## Movie legends

**Movie 1.**

Mesh-like F-actin networks cross-linked by passive ACP, anillin, are formed in the absence of myosin. This particular network is comprised of 3 μM G-actin, 0.90 μM anillin, and 1 mM ATP. Rhodamine phalloidin labeled-F-actin molecules are shown. Time is given as h:min:s. Scale bar, 100 μm.

**Movie 2.**

Actomyosin forms small clusters with the mean maximum cluster size of 100 μm^2^ in the absence of anillin. This particular network is comprised of 3 μM G-actin, 1 μM myosin minifilaments, and 1 mM ATP. Rhodamine phalloidin-labeled F-actin molecules are shown. Time is given as h:min:s. Scale bar, 100 μm.

**Movie 3.**

Actomyosin forms medium clusters with the mean maximum cluster size of 100 to 1,000 μm^2^. This particular network is composed of 3 μM G-actin, 1 μM myosin minifilaments, 0.36 μM anillin and 1 mM ATP. Rhodamine phalloidin-labeled F-actin molecules are shown. Time is given as h:min:s. Scale bar, 100 μm.

**Movie 4.**

Actomyosin forms large clusters with the mean maximum cluster size over 1,000 μm^2^. This particular network is composed of 3 μM G-actin, 1 μM myosin minifilaments, 0.72 μM anillin and 1,000 μM ATP. Rhodamine phalloidin-labeled F-actin molecules are shown. Time is given as h:min:s. Scale bar, 100 μm.

**Movie 5.**

Actomyosin does not form clusters at a still higher anillin concentration. This particular network is composed of 3 μM G-actin, 1 μM myosin minifilaments, 0.90 μM anillin and 1 mM ATP. Rhodamine phalloidin-labeled F-actin molecules are shown. Time is given as h:min:s. Scale bar, 100 μm.

**Movie 6.**

Myosin minifilament-induced F-actin severing occurs in a 3D disordered network. The 3D disordered network is composed of 0.1% rhodamine phalloidin-labeled F-actin, 3 μM dark G-actin, 0.5 μM anillin, 1 μM myosin minifilaments, and 1 mM ATP. Rhodamine phalloidin-labeled F-actin molecules are shown. Time is given as in h:min:s. Scale bar, 5 μm.

**Movie 7.**

Actin severing protein, gelsolin, induced network contraction. The prestressed networks containing 3 μM G-actin, 0.72 μM anillin, 1 μM myosin and 50 μM ATP with ATP regenerating system were formed (10 s intervals, ×50 actual speed). At 360 sec, gelsolin (left, 10 s intervals, ×100 actual speed) or buffer solution (right, 10 s intervals, ×400 actual speed) was added into the prestressed network. Rhodamine phalloidin-labeled F-actin molecules are shown. Time is given as h:min:s. Scale bar, 100 μm.

